# Honey bee worker jelly, a nutritional secretion fed to offspring, shows high among-nestmate and among-colony variation

**DOI:** 10.1101/2025.06.26.661879

**Authors:** R. R. Westwick, C. C. Rittschof

## Abstract

Parents have evolved strategies to reduce the risk of malnutrition in offspring, including the production of specialized nutritional secretions that are tailored to meet the unique needs of developing offspring. Studies in vertebrates, however, show surprising individual variation in nutritional secretions; the causes and consequences of this variation, and the extent to which such patterns can be generalized beyond vertebrates, remain unclear. Here, we investigated natural variation in nutritional secretions in an invertebrate species, the honey bee (*Apis mellifera* L.). This is a unique bee wherein developing larvae subsist entirely on “jellies” (e.g., royal jelly) produced by glands in adult worker bees. We assess among nestmate and among-colony variation in the macronutrient content of the secretions fed to female worker larvae (“worker jelly”). Although female workers make up the largest demographic inside a honey bee colony, very few studies have investigated their larval diet; even fewer have included the scope of colonies needed to assess natural variation in this critical nutritional substance. In one of the largest such studies to date, we found significant variation both among nestmates and among colonies in total quantity and macronutrient content of worker jelly. This pattern was strongest, surprisingly, for proteins and lipids. Moreover, the macronutrient ratio in worker jelly differs substantially from source pollen, suggesting adult workers retain substantial consumed nutrients, especially lipids. We further assessed whether worker jelly composition was correlated with colony defensive aggression because of extensive links between aggression, foraging activity, and larval development outcomes; however, we observed no such relationship. This study is a critical step in understanding the evolution and maintenance of offspring provisioning strategies, as well as bee foraging ecology and nutritional stress response.

## INTRODUCTION

Early-life nutrition has long-lasting consequences for an organism’s growth and survival, as deficits during this high-growth time often cannot be fully compensated for later (1–3). Many organisms have evolved parental care to hedge against early-life risks; this often involves adults provisioning young to ensure adequate nutritional intake (4). Sometimes adults simply assist their offspring in finding naturally available nutritional resources (5). But in other cases, caregivers gather and process nutritional resources before providing them to the young. For example, some birds will regurgitate partially digested stomach contents for their chicks (6). In the most evolutionarily derived form of provisioning, adults have specialized body structures to synthesize and secrete nutritional products tailored for their young (such as the milk produced by mammals; (7)). Nutritional secretions occur in a wide variety of taxa, such as pigeons, bony fish, caecilian amphibians, and some social insects, among others (8–13). In cases where young feed exclusively on secretions, organisms face the challenge of synthesizing a product that will meet all of the changing nutritional needs of their offspring across different growth stages (14). Accordingly, the contents of nutritional secretions are often dynamic and age-specific (15, 16). Therefore parents, who can sometimes even break down and incorporate their own body tissues if food resources are limited, presumably provide reliable and uniform complete nutrition for offspring.

Despite evidence that nutritional secretions track species- and development-specific nutritional demands, strong inter-individual variation in secretion composition is common, at least in the limited cases where it has been examined (17–22). This pattern is somewhat surprising (15), as there are costs to variation in secretions; for example, variation in individual human milk oligosaccharides has been correlated with a range of negative health outcomes in infants (23). The incidence, causes, and consequences of individual variation in nutritional secretions remain unclear.

Studies of variation in nutritional secretions have been conducted almost exclusively in mammalian systems, where one individual (the mother) is typically responsible for all the nutritional resources an offspring receives. Some social species, notably social insects including the honey bee (*Apis mellifera* L.), distribute brood care among many hundreds or even thousands of adult relatives (24–26). There may be adaptive benefits to both uniformity and diversity in nutrition. If one offspring is exposed to many caregivers, it may average out variation in total nutrition received, resulting in consistent and optimal offspring nutrition within a social group. Conversely, variation in resource availability to a colony—coupled with care from a variety of good and poor providers—could increase variation in the nutrition profiles for individuals within and among social groups.

Selection pressures could also shape realized variation in offspring nutrition. For example, if there is a species-specific optimal intake target, variation within and among social groups should be reduced where possible (27, 28). However, in some contexts, variation in offspring nutrition may be adaptive; in honey bees, adult worker diversity in phenotypes like cue response thresholds and daily activity patterns improve colony performance (29–35). While some of this diversity is the result of genetic variation among workers within a colony (36), the developmental environment also shapes these phenotypes (37–41). The role of developmental nutrition in these processes is virtually unstudied. For example, in free-living honey bees showing a spectrum of defensive aggression, cross-fostering studies show that the larval developmental environment shapes adult aggressive behavior and several other co-varying phenotypes (40, 41). Though these studies did not explicitly look at the role of nutrition within the context of the developmental environment, other studies have shown that caregiver interactions (including feeding visits) are different between relatively aggressive and docile colony environments (42). Together, these results suggest a role for variation in larval nutrition that remains untested. This notion is especially intriguing given the many links between nutrition and aggression that have been seen in other taxa (43–46).

In the current study, we measured the total amount and macronutrient content of the secretions (“worker jelly”) provided to over 450 individual workers from 18 colonies that show natural variation in defensive aggression. This is an unprecedented scope of colony-level sampling for honey bee larval nutrition; this study therefore represents a critical step towards determining the relationships between developmental nutrition and adult worker phenotypes, as well as the colony-level implications for such patterns. We hypothesized low natural variation in worker jelly macronutrients within colonies, especially proteins and lipids that are most critical to larval development (47), because nurses can use their own metabolic reserves to buffer natural fluctuations in colony resource conditions (48–50). We additionally hypothesized that high- and low-aggression colonies would show differences in the nutritional makeup of the worker jelly (either in terms of total food available or in terms of macronutrient ratios) because of the known relationship between aggressive social environments and the health and aggressive outcomes for larvae raised in them (40–42). In contrast to our predictions, we found high levels of variation in worker jelly nutrition both among nestmates within a single colony and among colonies, and this variation was not associated with aggression.

## METHODS

### Study system

Honey bee larvae are fed by a specialized group of adult workers called nurses, which consume pollen and honey and synthesize worker jelly using hypopharyngeal glands in the head (51). Worker jelly secretions are passed orally into honeycomb cells containing developing larvae. The larva typically sits in a pool of worker jelly which she ingests over time. Previous studies have determined the basic components of worker jelly (52, 53) and that the macronutrient profile and other nutritional contents change with larval age (52, 54), suggesting precise regulation of nutrition. Several hundred to several thousand nurse bees will be active at any given time, walking around the brood nest, checking on larvae, and feeding them if necessary (24). Nurses care for larvae around the clock, with a single larva being visited upwards of 3,000 times in a single day by the final developmental stage (55).

### Colony choice and sample collection

We collected worker jelly from 18 managed but otherwise unmanipulated honey bee colonies in central Kentucky, USA from June 25-July 21, 2019. Most colonies (N=10) originated from packages in late April with strains advertised as “Italian” and “Russian Hybrid” (Schoolhouse Bees, Covington, KY). Two colonies were installed as packages in May from two separate providers (Dadant & Sons, Frankfort, KY; Hosey Honey, Midway, KY) and no strain information was provided. Remaining colonies (N=6) were of local open-mated stock.

Colonies were kept at one of three sites near Lexington, KY, USA. All sites were on mixed-use farms with similar surrounding landscapes. Two of the sites were approximately 1.6 km apart (“Alpha” and “Beta”), which is within the maximum honey bee foraging range but beyond the typical range of most foraging trips (56–59). A third site (“Gamma”) was approximately 14.5 km away from the other two, which is outside of the maximum foraging range for Alpha and Beta (60). Variation in the surrounding landscape could impact the floral resources available to colonies (and thus worker jelly content); though we did not perform any direct assessments of landscape floral resources in this study, we do consider location as an explanatory variable in our analyses.

Because previous studies associated colony defensive aggression (hereafter aggression) with nurse care behaviors (42) and larval developmental outcomes (40, 41), we sampled worker jelly from colonies ranging in aggression phenotype. We assayed 36 colonies using a previously established aggression assay (Alaux & Robinson, 2007; Rittschof et al. 2015). Briefly, we took photographs at the entrance of the colony and counted the number of visible bees. We then placed a strip of filter paper containing 3 µL of 1:10 isopentyl acetate : mineral oil in the center of the entrance. Isopentyl acetate is the main active component of the honey bee alarm pheromone (61), and it provokes workers to emerge from the colony entrance; a greater number of emerging bees signifies higher aggression (62). One minute following the filter paper application, we took a second photograph to quantify the emergence response. We calculated an aggression score as the difference between the number of bees at the entrance after versus before filter paper application. All colonies at a given site were assayed for aggression on the same day. The 9 highest-aggression and 9 lowest-aggression colonies were chosen for worker jelly collections, blocked across the three sites (N=18 total colonies, N=3 high-aggression and N=3 low-aggression at each site). Each colony thus has a quantitative aggression score, a relative rank compared to the other 17 colonies in the study, and a categorization as high versus low aggression; all measures were incorporated in our data analyses (see below). We performed worker jelly collections at variable times but within 4-16 days following the aggression assays. This short experimental timeframe (26 total days to identify colonies and collect all samples) minimizes temporal variation in colony and climactic conditions that could influence worker jelly content and/or colony aggression score (63–67).

The macronutrient content of the worker jelly changes with larval age (52). We therefore sampled from age-matched larvae. Honey bee larvae develop within their own individual honeycomb cells into which nurse bees deposit worker jelly. Trapping the queen against appropriately sized honeycomb for a short period of time is a common strategy to collect age-matched worker offspring (68, 69). For each of the 18 colonies, we caged the mother queen with empty honeycomb for 24 hours to allow her to lay worker-destined eggs. The cage fit one standard wooden “deep” frame with worker-brood-sized honeycomb from a Langstroth beehive (∼48 cm x 3 cm x 23 cm). Two sides of the cage were covered by a “queen excluder,” a plastic grate that was large enough to allow adult workers to pass in and out freely and tend to young while being small enough to contain the larger queen (70, 71). After 24 h, the queen was released back into the colony, and the frame was placed back in the colony within the cage to prevent further laying by the queen. This approach generates dozens to hundreds of eggs that range in age from 0 to 24 h old.

Nurses begin making feeding visits to deposit worker jelly almost immediately once a larva hatches, which takes about three days following egg laying (55, 72). Two days after the eggs hatched (96-120 hours post-laying), we removed frames to collect worker jelly on a single day per colony between 9:00 and 11:00. Samples from all 18 colonies were collected on one of 8 sampling days (S10 Table). We brushed off adult bees and covered the frame with damp paper towels to prevent larva and worker jelly desiccation. We carried the frame into a building to collect the worker jelly (<5 min following frame removal). We haphazardly chose approximately 27 cells covering the entire laying area on one side of each frame, excluding any cells where the larvae were markedly large or small or had any abnormalities. We collected 467 total samples across the 18 colonies; some colonies (N=4) had fewer than 27 samples due to an insufficient number of usable larval cells on one side of the frame; this was usually the result of poor laying success by the queen.

Honey bee larvae sit in the base of honeycomb cells in a small pool of worker jelly. To collect the worker jelly, we pipetted 100 uL of deionized water into the cell which caused the larva to float to the top. We carefully removed the larva with a small metal queen grafting tool and placed it in a 1.5 mL microcentrifuge tube (Thomas Scientific), which we stored at −80⁰C. We then pipetted an additional 100 uL of deionized water into the cell (200 uL total). We drew the slurry of water and worker jelly in and out of the pipette several times to mix it and to loosen the jelly from the sides of the cell. We pipetted the mixture into another 1.5 mL microcentrifuge tube (Thomas Scientific), where it was stored at −80⁰C until chemical processing.

### Sample processing

#### Total wet and dry mass

Worker jelly has a gelatinous texture and is comprised of water, proteins, lipids, carbohydrates, and other material such as micronutrients and nondigestible material like fiber. To homogenize the worker jelly samples and break up clumps, we sonicated the samples for 5 min on 100% power with a Misonix S-4000 cup horn sonicator (Misonix, Newton, CT). We then determined the wet and dry mass of a 20 uL aliquot from each worker jelly sample and used these values to calculate total wet and dry mass for each sample.

#### Protein quantitation

We used a bicinchoninic acid (BCA) assay to quantify total protein mass per sample (Pierce™ BCA Protein Assay Kit, 23227, Thermo Fisher Scientific, Waltham, MA, USA). The assay was carried out according to the manufacturer’s instructions for a microplate preparation, with Bovine serum albumin (2 µg/µL) as the standard. The standard curve included volumes of 0 µL, 2.5 µL, 5 µL, 7.5 µL, 10 µL, and 20 µL of the standard. We measured protein from a sample fraction unique from the fraction used for the lipid and carbohydrate extractions (see below). We adjusted the volume of the starting fraction for each sample based on the sample’s total dry mass to ensure the protein readings fell within the assay’s standard curve; we accounted for this variation in calculations of total protein mass per sample (mg). We measured all samples in triplicate. Samples were randomly distributed across assay plates to avoid any technical bias caused by plate effects, though an individual sample’s triplicates were all on the same plate. The final color is stable for approximately half an hour; it was during this half-hour window that we measured the absorbance on a microplate reader (CLARIOstar Plus, BMG LABTECH, Offenburg, Germany). We used the median triplicate value of each sample to calculate the total protein mass per sample (mg). We also divided total protein mass by total dry mass to calculate the relative protein quantity per sample (a unitless value).

#### Lipid and carbohydrate quantitation

Lipid and carbohydrate levels were measured from a single 100 µL aliquot on which we performed combined chloroform-methanol extraction and fractioning (following (73)). We performed a sulfo-phospho-vanillin assay on the chloroform fraction to determine the amount of lipids per sample. Commercial vegetable oil suspended in chloroform (1 µg/µL) was used as the standard, with volumes of 0 µL, 12.5 µL, 25 µL, 50 µL, and 100 µL for the standard curve. An anthrone-sulfuric acid assay was used on the methanol fraction to determine carbohydrate concentrations. Anhydrous glucose dissolved in deionized water (1 µg/µL) was used as the standard for this assay, with the same range of volumes for the standard curve. As with the protein assays, we adjusted the volume of the starting fraction for each sample based on the sample’s total dry mass to ensure the readings fell within the assay’s standard curve; we accounted for this variation in calculations of total lipid and carbohydrate mass per sample (mg). Samples were once again run in a random order. We also measured these samples in triplicate, although due to the nature of the assay, the samples were not split into three replicates until after the color change had occurred. Similarly to the protein assays, we measured the absorbance on a CLARIOstar microplate reader during the half-hour window that the color change is stable. The median value of each triplicate was used to calculate total lipid mass per sample (mg) and total carbohydrate mass per sample (mg), as well as relative lipid and carbohydrate quantities (unitless) as described above for protein.

### Statistical analyses

We performed statistical analyses with R version 4.1.2 (74). Our analyses primarily focused on total dry mass (mg); total protein, lipid, and carbohydrate mass (mg); and relative protein, lipid, and carbohydrate quantities (unitless) as response variables. To run a Principal Components Analysis (PCA) on macronutrient masses and relative quantities, we used the prcomp() function in the “stats” package (74). The “ggbiplot” package was used to create visualizations of the PCA (75).

We used the “lme4” package in R to create separate linear mixed models (LMMs) to assess the fixed effects aggression, colony site, and/or colony genetic strain separately for each response variable (macronutrient masses and relative quantities) (76). We incorporated aggression into the analyses in three ways (with separate models for each): as a binary variable (high/low), as a ranked variable (1–18), and as a continuous variable using quantitative aggression scores. Colony site and colony genetic strain each had three levels (Alpha, Beta, or Gamma; and “Italian,” “Russian hybrid,” or open-mated). Square-root and log+1 transformations were used as needed to normalize the data and improve model fit. Total dry mass was either square root or log+1 transformed depending on the analysis (stated individually in the RESULTS). Unless otherwise stated, protein, lipid, and carbohydrate total masses and relative quantities were square-root transformed. Our initial models each contained a single fixed effect (aggression level, aggression rank, quantitative aggression score, colony site, or genetic strain) and colony identity as a random effect. We additionally created one model that included multiple fixed effects: aggression, colony genetic strain, and their interaction, with colony ID as a random effect. We used the DHARMa package to assess the quality of model fit, which included a Q-Q plot, Kolmogorov–Smirnov test, dispersion test, outlier test, group-level uniformity test, and Levene test (77). We used Type II ANOVAs to assess significance values for our models; for this we used the “car” package (78). To calculate among-colony variance, we used the “aov” function from the “stats” package to run ANOVAs (74). To perform median-centered Levene’s tests for homogeneity of variance, we used the “leveneTest” function from the “car” package (78). Figures were created using the “ggplot2,” “ggpubr,” “cowplot,” and “viridis” packages (79–82).

## RESULTS

### General observations

We measured macronutrients in 467 samples of worker jelly from 18 different colonies. Our samples had an average total dry mass of 1.24 ± 0.63 mg (mean ± SD) with 0.82 ± 0.50 mg of total proteins, 0.05 ± 0.03 mg of total lipids, 0.10 ± 0.06 mg of total carbohydrates, and 0.26 ± 0.34 mg of other matter (e.g. nondigestible material, micronutrients, minerals, contaminants, etc.). These measurements give an average ratio of approximately 16 : 1 : 2 : 5 proteins : lipids : carbohydrates : other matter, or 66.6% proteins, 4.4% lipids, 7.9% carbohydrates, and 21.1% other matter. S11 Table shows how these values compare with previous studies that measured worker jelly composition. Although there is considerable variation in worker jelly nutrient composition among studies, our measurements are within the ranges reported by others (e.g., (52, 83, 84)) and our results are in close agreement with Wang et al. (2015), the most recent study that examined a similar-aged cohort of larvae.

We assessed the major factors driving relationships among samples using a Principal Components Analysis (PCA). We performed two analyses, one using macronutrient masses and one using relative macronutrient quantities (omitting the “other” category; Fig 1). For mass, the first component (PC1), which was largely attributable to variation in protein, explained 75.6% of the total variance. The second component (PC2) explained 15.9% of the variance for a total of 91.5%. PC2 depicted an inverse relationship between lipids and carbohydrates. The PCA for relative macronutrient quantities gave similar results, but with slightly less total variance explained (88.5%), and greater evidence of covariation between lipids and proteins. Fig S1 displays these PCAs as a function of aggression level; there was very little separation between groups (see more detailed analyses of aggression below).

**Fig 1.**
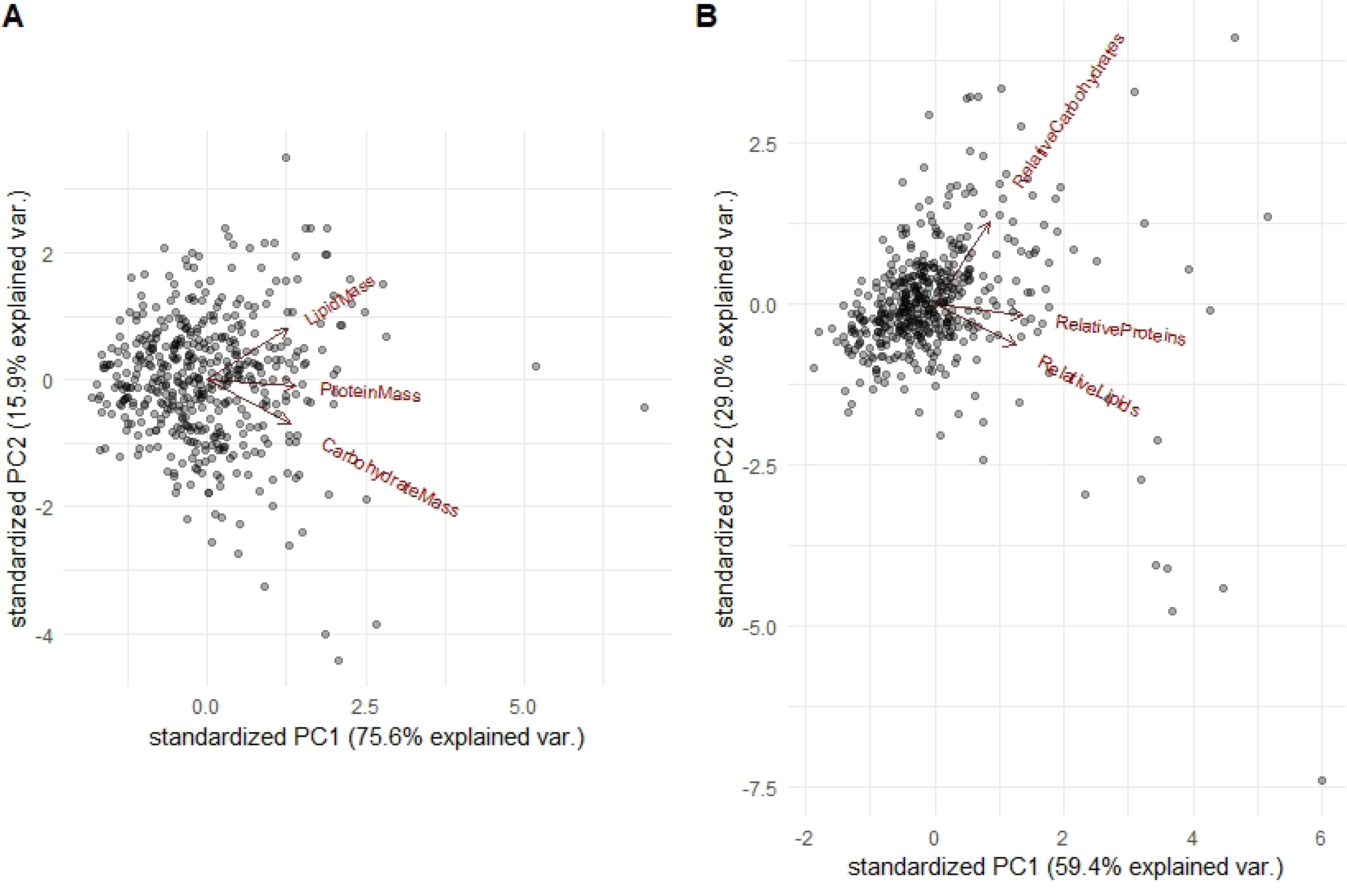
Principal Components Analysis of (A) total protein, lipid, and carbohydrate mass (mg) as well as B) relative quantities. PC1 for both is largely driven by protein quantity, while PC2 loads lipids and carbohydrates opposite of one another.

### Among-nestmate and among-colony variation in worker jelly quantity and content

There was considerable variation among colonies in the mean total dry mass per larval sample; colony means ranged from 0.4-2.1 mg (Fig 2, A). Mean total mass of each macronutrient also varied substantially among colonies (proteins: 0.27-1.49 mg, lipids: 0.02-0.11 mg, carbohydrates: 0.03-0.16 mg), as did relative macronutrient quantities to a lesser degree (Fig 2, B). Though we focus our discussion on relative macronutrient quantities in later sections of the results, here we focus on the total macronutrient masses to more intuitively discuss the magnitude of variance (notably, relative quantities gave similar statistical results to total macronutrient masses for all analyses). Variance among colonies was higher than variance among nestmates within a single colony for total dry mass and total mass of each macronutrient (ANOVA: total dry mass (square-root transformed): F_17_=24.4, P<0.0001; total protein mass: F_17_=26.1, P<0.0001; total lipid mass: F_17_=17.3, P<0.0001; total carbohydrate mass: F_17_=20.1, P<0.0001). Total protein mass showed the most extreme differences among colonies relative to the differences among nestmates, and total lipid mass showed the smallest differences among colonies relative to differences among nestmates (although still significant colony-level variation); total carbohydrate mass showed patterns in between proteins and lipids. Additionally, total protein mass showed significant non-homogeneity of variance among colonies, meaning that some colonies showed significantly higher variation among nestmates than others (Levene’s test: total protein mass: F_17_=2.76, P=0.0002). No other macronutrient showed this pattern (Levene’s test: total lipid mass: F_17_=1.36, P=0.15; total carbohydrate mass: F_17_=0.91, p=0.56), and neither did total dry mass (Levene’s test: total dry mass square-root transformed: F_17_=0.88, P=0.60). Overall, protein appears to be the macronutrient that has both the greatest variation among colonies and the greatest range of variances among nestmates.

**Fig 2.**
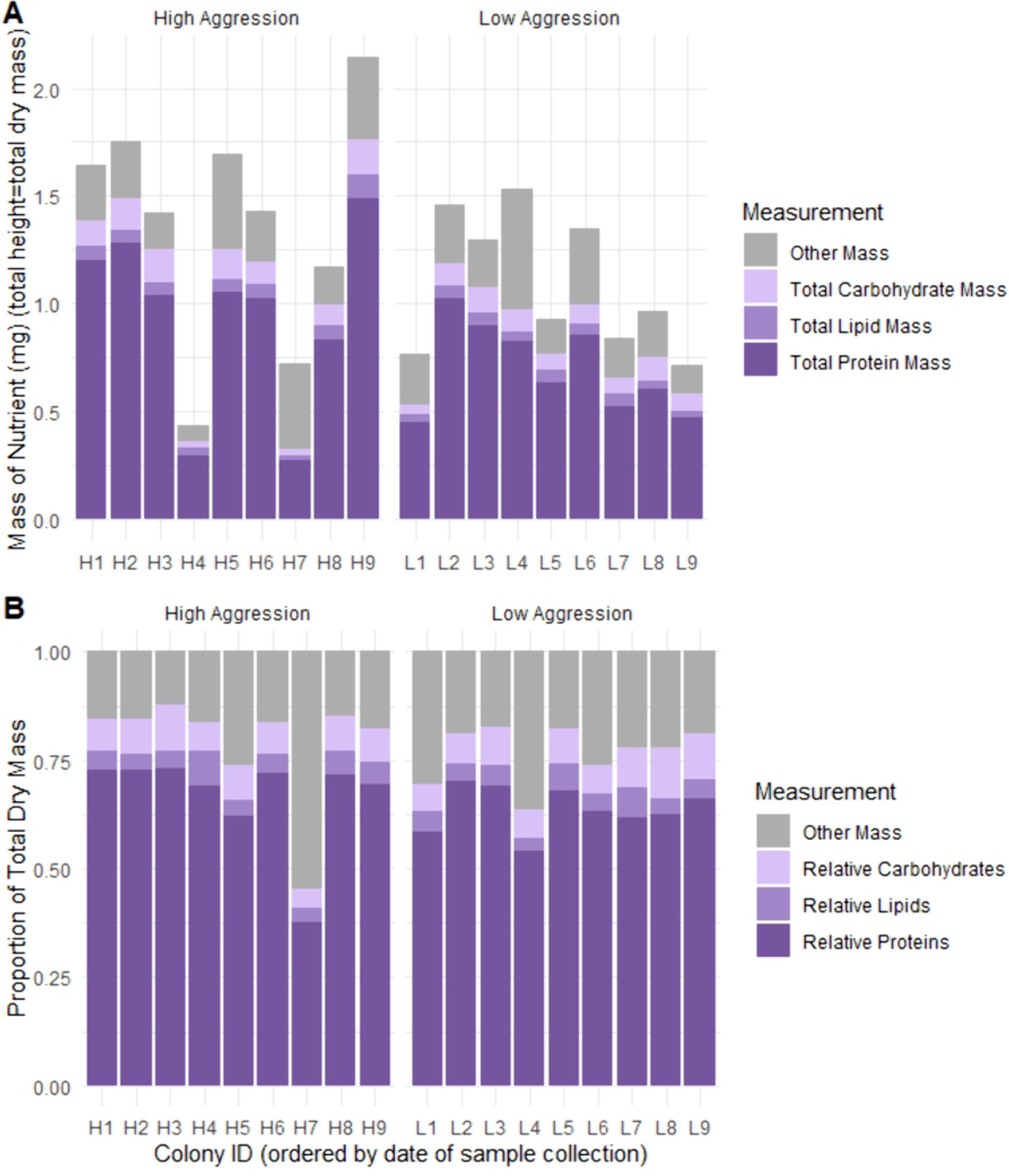
Colony variation in total dry mass, and total and relative mass of each macronutrient. High-aggression colonies (H) are clustered on the left and low-aggression colonies (L) are clustered on the right in each panel. A) Per-colony mean total mass per sample of each macronutrient (mg). The total height of each bar is the mean total dry mass per sample (mg). B) Per-colony mean proportion of each macronutrient relative to the total dry mass of each sample (which is set at 1).

### Relationship between colony aggression and worker jelly quantity and content

Aggression scores for all 18 colonies are shown in Fig S2. To assess the relationship between colony aggression and worker jelly quantity and content, we first treated aggression as a binomial variable (low versus high) in LMMs. This analysis showed no association between colony aggression and total dry mass or relative macronutrients quantities (Fig 3; total dry mass (log-transformed): Wald X^2^_1_=1.3, P=0.25; relative protein quantity: Wald X^2^_1_=0.64, p= 0.42; relative lipid quantity: Wald X^2^_1_=0.07, P= 0.79; relative carbohydrate quantity: Wald X^2^_1_=0.42, P=0.51). Similar models with total macronutrient masses (mg) gave comparable results (Fig S3), as did analyses that treated aggression as either a ranked variable or a continuous variable (1-18, see METHODS; Fig S4-S6; (40)).

**Fig 3.**
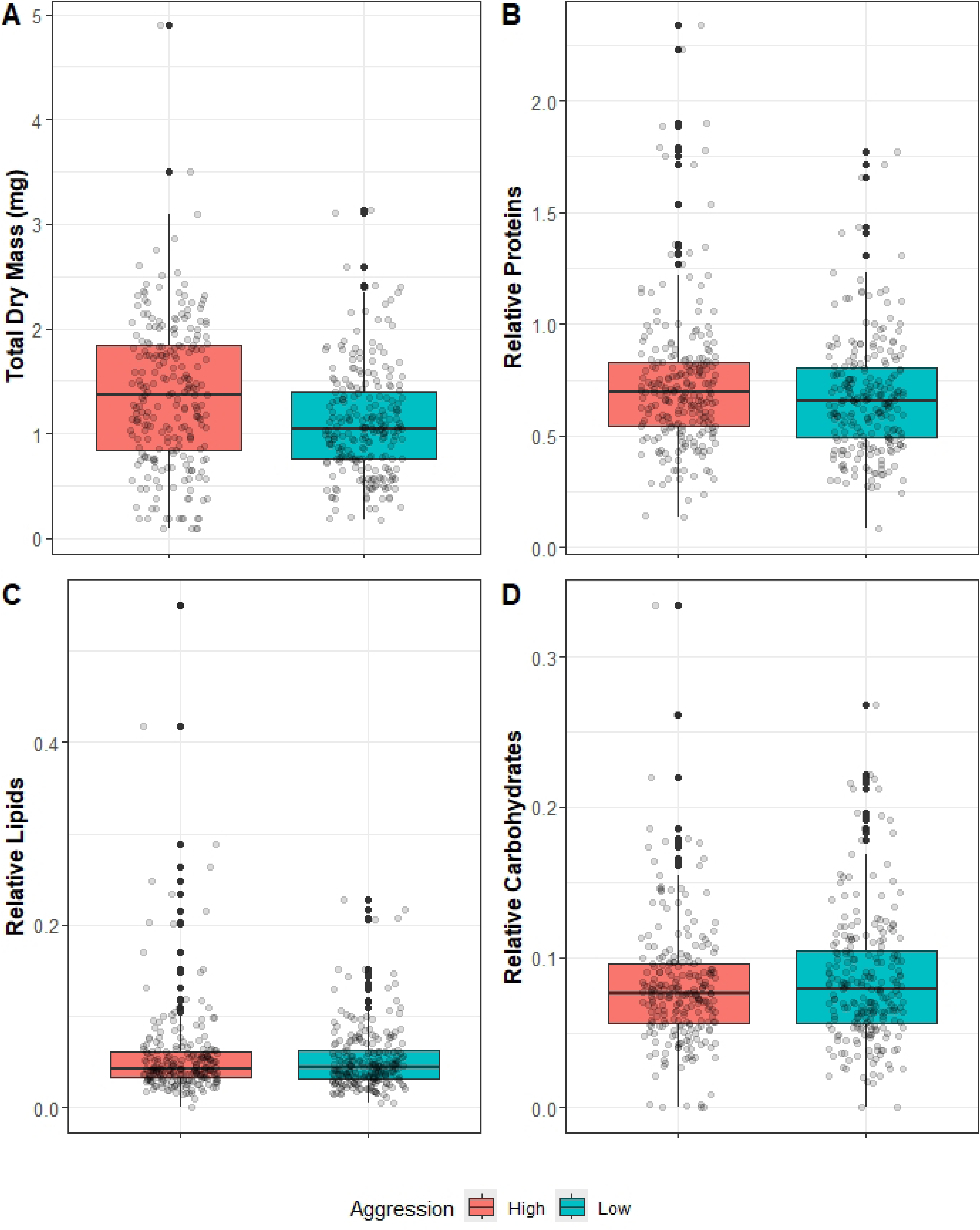
Total dry mass and relative amounts of each macronutrient in worker jelly as a function of colony aggression level (High versus Low). Boxplots show the distribution (1st quartile, median, and 3rd quartile for the box and 1.5x the IQR for the whiskers) for A) total dry mass (mg), B) relative proteins, C) relative lipids, and D) relative carbohydrates. Each dot represents a unique worker jelly sample. Fig S5 shows panels for total proteins, total lipids, and total carbohydrates (all mg).

Because the high aggression colonies included some of the highest and lowest per-colony mean total dry masses (Fig 2), we assessed how variance in worker jelly differed between high-and low-aggression colonies. Like the whole-data analyses described above, we used ANOVAs for total dry mass and relative macronutrient quantities within each aggression level to test how among-nestmate versus among-colony variance changed with aggression level. The difference between among-colony and among-nestmate variance (the magnitude of the F-statistic) was much higher in high-aggression colonies for all measures, although we identified significant colony-level variation in each macronutrient across both aggression levels (ANOVA: total dry mass (square root-transformed): high: F_1_=33.6, P<0.0001; low: F_1_=13.5, P<0.0001; relative protein quantity: high: F_1_=38.1, P<0.0001; low: F_1_=10.4, P<0.0001; relative lipid quantity: high: F_1_=28.4, P<0.0001; low: F_1_=6.5, P<0.0001; relative carbohydrate quantity: high: F_1_ =33.5, P<0.0001; low: F_1_ =7.5, P <0.0001).

### Alternative hypotheses to explain variation in worker jelly content

We designed our experiment to assess whether colony aggression level explained variation in worker jelly content, but we performed some additional analyses to assess other possible contributing factors. We first assessed evidence of temporal variation over the course of our experimental timeframe. An ANOVA across the 8 sample collection days (see METHODS) showed that collection date did not explain significant variation in total dry mass or mass of any macronutrients (Fig S7; total dry mass (square root-transformed): Wald X^2^_1_=0.81, P=0.37; relative protein quantity: Wald X^2^_1_=0.25, P=0.61; relative lipid quantity: Wald X^2^_1_=0.33, P=0.56; relative carbohydrate quantity: Wald X^2^_1_=0.42, P=0.52).

Next, we assessed evidence of site-level variation in worker jelly. We found that site was not associated with significant variation in in any of our measures (LMMs with site as a fixed effect and Colony ID as a random effect; total dry mass (log-transformed): Wald X^2^_2_=2.9, P=0.24; relative proteins: Wald X^2^_2_=0.7, P=0.70; relative lipids: Wald X^2^_2_=2.6, P=0.28; relative carbohydrates: Wald X^2^_2_=1.4, P=0.50, Fig S8). Results were similar for total macronutrient masses (data not shown).

Finally, we examined effects of colony genetic background on worker jelly content, focusing on 411 samples from 16 colonies representing three genetic “strains”, those advertised as “Italian” (N=4) or “Russian Hybrid” (N=6), and “Open-mated” colonies from a local beekeeper (N=6) (85). Two additional strains (each with N=1 colonies) were excluded due to low replication (see METHODS). We found no evidence of significant variation in worker jelly total dry mass or relative macronutrient quantities as a function of genetic strain (LMM, total dry mass (log-transformed): Wald X^2^_2_=5.6, P=0.06; proteins: Wald X^2^_2_=0.9, P=0.64; lipids: Wald X^2^_2_=2.5, P=0.29; carbohydrates: Wald X^2^_2_=2.5, P= 0.29, Fig S9). Results were similar for total macronutrient masses (data not shown). Because aggression is partially heritable (86–88), we assessed evidence for an interaction between aggression level and strain. We did not detect statistically impactful collinearity between aggression and genetic strain (generalized variance inflation factor (GVIF)—all values below 2). We therefore ran an additional model that included both genetic strain and aggression as well as an interaction term between aggression and genetic strain, but this model yielded similar nonsignificant results (data not shown).

## DISCUSSION

In one of the largest modern studies of honey bee worker jelly content (over 450 samples from 18 unique colonies), we observed substantial and significant variation even for age-matched samples, both among colonies and among nestmates within a single colony; notably, this variation existed despite our measurements of total quantity and macronutrient composition falling generally within the ranges of previous studies. Proteins, lipids, and carbohydrates all showed significant among-colony variation. Protein is the greatest proportion of worker jelly, and it is often discussed as one of the most critical components of insect larval development (89–92); proteins not only showed the greatest among-colony variation, but colonies also differed substantially in the extent to which samples from different nestmates varied in this nutrient. Lipids, the smallest worker jelly component, showed the smallest differences among colonies relative to differences among nestmates (although still significant colony-level variation). Carbohydrates, arguably the most easily acquired macronutrient due to its presence in both pollen and nectar/honey, showed an intermediate level of among-colony variation relative to among-nestmate variation (47, 93). In terms of explanations for among-colony variation in samples, we found little evidence that colony defensive aggression played a clear role, although the magnitude of among-colony variation relative to among-nestmate variation was higher in high aggression colonies. No other measured factor (sample collection date, site, colony genetic strain) explained variation in worker jelly composition.

Surprisingly, even with multiple caregivers, the worker jelly nutritional secretion fed to age-matched larvae within a single colony showed high variation, similar to what is seen in species with a single maternal caregiver, e.g., mammals (17–19). However, despite this relatively high among-nestmate variance, among-colony variance was significantly higher, indicating that a complex combination of factors likely impacts the nutritional experience of individual worker honey bee larvae. Variation in lipid and protein content is surprising given their critical role in larval development. One possibility is that, once a minimum macronutrient threshold is met, any further variation is less important for health and fitness consequences. Natural selection would drive honey bees to meet a minimum level of each macronutrient, but quantities exceeding these levels would not necessarily be selected against (94). Honey bee larvae could also represent what is referred to in nutritional ecology as “macronutrient generalist”—that is, a species that is able to physiologically tolerate a wide range of macronutrient ratios (95). Overall, though recent research has suggested that nurse bees may regulate their dietary intake of protein-to-lipid ratios (96), our work suggests that this regulation does not lead to stable protein-to-lipid ratios in the worker jelly that these nurses produce as much as might be expected, at least from the perspective of the larvae that rely on these nutrients. However, few other studies have extensively sampled macronutrients from worker jelly; data on lipids is particularly scarce (S11 Table). More studies are thus required to understand the nutritional needs of honey bee larvae.

We found that the ratio of proteins to lipids in worker jelly is about 16:1, which is more protein-biased than most measurements of the royal jelly that is produced for rearing queens (ranging from 3:1 to 16:1 in different studies; e.g. (97–99)), and remarkably higher than ratios identified in honey bee-collected pollens (1.5:1) (100). Collectively, these results suggest an outsized role for nurse bee nutrient processing in larval dietary macronutrients, and an overall loss of lipids comparing source pollen to processed food. In previous research, nurse bees either do not seem to show nutrition-based preferences for particular pollens or show distinct preferences from foragers, which could contribute some to the difference between forager input and nurse output (101–103). However, nurse bee physiological needs may also play an important role in this discrepancy. In the first two weeks following adult emergence, worker honey bees increase their stored fat, the majority of which accrues in the multipurpose fat body organ (104). This lipid assimilation corresponds to the period when adult workers specialize on nursing behaviors (105), and lipid consumption during the nursing timeframe is important for development and maintenance of the hypopharyngeal glands used to generate worker jelly (106–110). A significant amount of the lipids that nurses consume appear to be retained rather than converted into worker jelly, which suggests they do not typically contribute lipids from their own fat stores to worker jelly under normal nutritional conditions (106, 111, 112). This could explain why older workers are able to perform nursing behaviors even with very low fat stores (113, 114). Proteins could show opposite patterns, however, since they show enhanced concentration in worker jelly relative to source pollen. Indeed, nurse bees that are well-fed show enhanced expression of protein-secretion-related genes in their hypopharyngeal glands, suggesting that endogenous protein production is increased even in the presence of ample food (115); additionally, the social pheromone QMP seems to mobilize abdominal protein for the purposes of larval jelly production (116). Further work is needed to understand the potential for nurse bees to protect larvae from variation in floral resource availability—this ability may be nutrient-specific.

Although nurse bee nutrient acquisition could explain the disconnect between source pollen and worker jelly macronutrient content, it does not explain the variability among worker jelly samples, nor why patterns of variation differ across macronutrients. Proteins and lipids largely come from the same source (pollen) whereas carbohydrates come primarily from nectar and/or honey (47). As a result, floral diversity and pollen source are expected to have similar impacts on lipid and protein content compared to carbohydrates, a pattern supported in some studies (117, 118). Although our results reflect these dynamics somewhat—for example, our principal component analysis showed that proteins and lipids were weighted more closely to one another than to carbohydrates – we still find distinct patterns of variation for each macronutrient, and that pattern remains unexplained.

We designed our study to assess whether variation in colony aggression level predicted the quantity or make up of worker jelly. We based our sampling design and scale on previous studies where a similar approach found significant effects of colony aggression on adult defensive aggression and immune system function for workers developing in these environments (40, 41); aggression-related variation in nursing behavior suggests nutrition could play a key role in these patterns (42). In all of these studies, including the current study, variation in aggression likely originated from a combination of genetic and environmental sources. While aggression—and associated variation in adult worker behavior and physiology (40, 41, 119–122)—did not explain significant variation in worker jelly quantity and content, among-colony variance was greater for high-aggression colonies than low-aggression colonies. This could suggest that diverse factors can drive up aggression levels, only some of which are related to brood provisioning. Variation in nursing behaviors might suggest that high aggression colonies have variable or less predictable food availability over time (42, 123), a characteristic that can engender physiological resilience later in life (39, 124) and may explain some positive health outcomes associated with aggression. Alternatively, among-nestmate variance was relatively more important in low-aggression colonies compared to high-aggression colonies. Developmental environments characterized by low aggression have been associated with a stressed immune phenotype in emerging adult honey bees (41); perhaps the low-aggression colonies struggle to hit nutritional targets as consistently as their high-aggression counterparts, where nutrients may frequently be in excess.

As with many colony-level honey bee studies, we noted several unusual colonies in our data. For example, the highest colony-average total dry mass per sample (found in colony H9) is nearly five times higher than the lowest colony-average total dry mass per sample (colony H4). Additionally, one colony (H7) had a significantly higher proportion of mass in the “other” category—that is, mass that is not accounted for by proteins, lipids, or carbohydrates—than any other colony in this study. This latter phenomenon would likely not be due to technical error in the sample collection or measurement process, as all of H7’s samples were collected and weighed at the same time as another colony, L7, which showed typical amounts of “other” mass, and all samples were randomized between plates during the colorimetric assays. We do not have a definitive explanation for this result, nor do we know what comprises the “other” mass, though it could be anything from nondigestible fiber to contaminants (125–129). This study was limited to three major macronutrients: proteins, lipids, and carbohydrates. Many other components are known to be present in worker jelly, such as phytochemicals, vitamins, minerals, other micronutrients, microorganisms, and bioactive compounds such as neurotransmitters and hormones (47, 53, 54, 130). Future work could assess the intra- and inter-colony variability of these smaller components, which may help explain some of these abnormalities noted in our experiment.

Our study took the larval perspective and examined the food in a cell available to a larva at a single point in time, which could include additions from multiple nurses. Future studies could seek to determine the level of individual variation from the perspective of nurse bees by extracting worker jelly from individual nurses and comparing it with that of their colony mates. Similarly, our study—which focused on 2-day-old larvae—cannot say whether other larval stages would show more, less, or a similar amount of variation associated with measured variables.

Additionally, we did not measure individual or colony-level outcomes for the larvae in our study. It is possible that the significant natural among-nestmate and among-colony variation we observed would not be enough to cause lethal or sublethal effects on the bees or colonies. However, potential outcomes based on previous experimental work are diverse and range in subtlety. Higher rates of developmental failure during the pupal stage and changes in body weight and morphometrics are associated with larval food deprivation (131, 132). Larvae from colonies with an experimentally reduced nursing workforce show a ∼25% decrease in adult lifespan and altered morphometrics relative to those with sufficient nursing effort (133, 134). Even minor underfeeding of larvae (much lower than the range of variation we observed in our study) can lead to reduced adult body size (135), and larvae artificially reared on diets with non-plant-based protein sources show relatively normal body weights and survival rates, but have altered behaviors and hormone profiles in adulthood (136, 137). Variation in carbohydrates (also observed in our study but to a lesser degree than proteins) in lab-based larval diets affects queen/worker caste determination (138). Thus, it seems likely that the degree of variation in worker jelly quantity and content we observed is sufficient to contribute to adult phenotypic variation.

Theoretical and experimental work suggests that parental care can buffer against environmental variability and risk on an evolutionary scale (139). The role of within-species variation in these dynamics, particularly in social species with cooperative brood care, provides an exciting new avenue for studying the developmental, physiological, behavioral, and health consequences of the early-life period.

## Acknowledgements

The authors would like to acknowledge and thank several people, without whom this experiment would not have been possible. First, we would like to thank Doug Potter for allowing us to use several of his honey bee colonies for this experiment, for helping with the field work on days we assayed and collected from his hives, and additionally for allowing us to use space near his apiary to harvest the worker jelly. We would also like to thank James Harrison and Anna Foose for their assistance with aspects of the field work, such as queen caging and retrieving the brood frames. Additionally, we would like to thank Jason Unrine for allowing us to use his sonicator and Jason Lichtenberg for training R.R.W. on its use. And finally, we would like to thank Joseph Palmer for his assistance in developing, optimizing, and training R.R.W. on the colorimetric assays used in this study.

## Ethics Statement

No approvals, licenses, or permissions were required to carry out this work. All honey bee colonies used in this study were maintained according to best practices recommended by the Honey Bee Health Coalition.

## Data and code

All data and code used to generate the results of this manuscript have been uploaded to Dryad (https://doi.org/10.5061/dryad.rjdfn2zpw),

Reviewer link: http://datadryad.org/share/ZXiQmy11J5c7um71L-XbowJmBTwkXQbrRgUA-VMaj7M

## Supporting Information

**S1 Fig. Principal Components Analysis (PCA) of A) total mass, and B) relative (“MassCorrected”) proteins, lipids, and carbohydrates with ellipses showing colony aggression level (high vs. low).** Note high degree of overlap between categories, with slightly more variation in high-aggression colonies along PC1 of A). PC1 of both figures is largely driven by protein measurements, while PC2 loads lipids and carbohydrates opposite of one another.

**S2 Fig. Quantitative aggression score as a function of binomial aggression level.** Boxplots show the distribution (following the convention in Figure 3) of raw aggression scores for colonies that were binned into the “high aggression” and “low aggression” categories; each dot represents one colony’s score. The raw aggression score was calculated as the difference between the number of bees at the colony entrance after an alarm pheromone presentation versus at baseline (see METHODS for details).

**S3 Fig. Colony-level aggression was not predictive of worker jelly nutritional content.** Linear mixed models of each nutrient with site as a fixed effect and colony ID as a random effect all showed no significant differences. Boxplots of A) total dry mass, B) total protein mass, C) relative proteins, D) total lipid mass, E) relative lipids, F) total carbohydrate mass, and G) relative carbohydrates of worker jelly samples for high-aggression (red) and low-aggression (blue) colonies.

**S4 Fig. Scatterplots of total dry mass (mg) and relative quantities of different macronutrients in worker jelly as a function of colony aggression rank.** Each dot represents an individual sample, black lines indicate the line of best fit (assuming a linear model), separated by aggression level. Grey shading around lines indicates the 95% confidence interval. Aggression rank (x-axis) is centered around zero, where “Low Aggression” colonies are negative numbers and “High Aggression” colonies are positive. Panels indicate A) total dry mass (mg), B) relative proteins, C) relative lipids, and D) relative carbohydrates.

**S5 Fig. Ranked aggression score is not predictive of worker jelly nutritional profiles.** Scatterplots of aggression ranks (centered around zero) versus A) total dry mass, B) total protein mass, C) total lipid mass, and D) total carbohydrate mass of honey bee worker jelly samples. Note the use of total masses of proteins, lipids, and carbohydrates in this figure compared to in-text figures; relative proteins, lipids, and carbohydrates were qualitatively similar. Colors are indicative of how colonies were grouped into the high-versus low-aggression categories in the previous analysis where aggression was treated as a binomial variable.

**S6 Fig. Continuous aggression score is not predictive of any of the nutrients we measured.** Scatterplots of raw aggression score versus A) total dry mass, B) total protein mass, C) total lipid mass, and D) total carbohydrate mass of honey bee worker jelly samples. Note the use of total masses of proteins, lipids, and carbohydrates in this figure compared to in-text figures; relative proteins, lipids, and carbohydrates were qualitatively similar. Colors are indicative of how colonies were grouped into high-versus low-aggression colonies in the previous analysis.

**S7 Fig. There is no clear seasonal trend over the course of the 23 days that samples were collected.** All charts show the average mass in mg per nutrient by experimental day (with the first collection day being Day 1), separated by colony aggression level. A) Total dry mass (mg), B) relative protein quantity, C) relative lipid quantity, D) relative carbohydrate quantity.

**S8 Fig. Total dry mass and relative quantities of each macronutrient (total nutrient mass divided by total dry mass for each sample) as a function of colony site.** Boxplots of A) total dry mass, B) relative proteins, C) relative lipids, and D) relative carbohydrates of worker jelly samples from three sites, Alpha (yellow), Beta (green), and Gamma (blue). Sites Alpha and Beta were approximately 1.6 km apart, while Gamma was approximately 14.5 km away from the other two sites. All comparisons are statistically nonsignificant based on linear mixed models.

**S9 Fig. Total dry mass and relative quantities of each macronutrient (total nutrient mass divided by total dry mass for each sample) as a function of colony site.** Boxplots of A) total dry mass, B) relative proteins, C) relative lipids, and D) relative carbohydrates of worker jelly samples from colonies headed by queens from three genetic strains, Italian (purple), Russian Hybrid (red), and open-mated (yellow). All comparisons are statistically nonsignificant based on linear mixed models.

**S10 Table.**
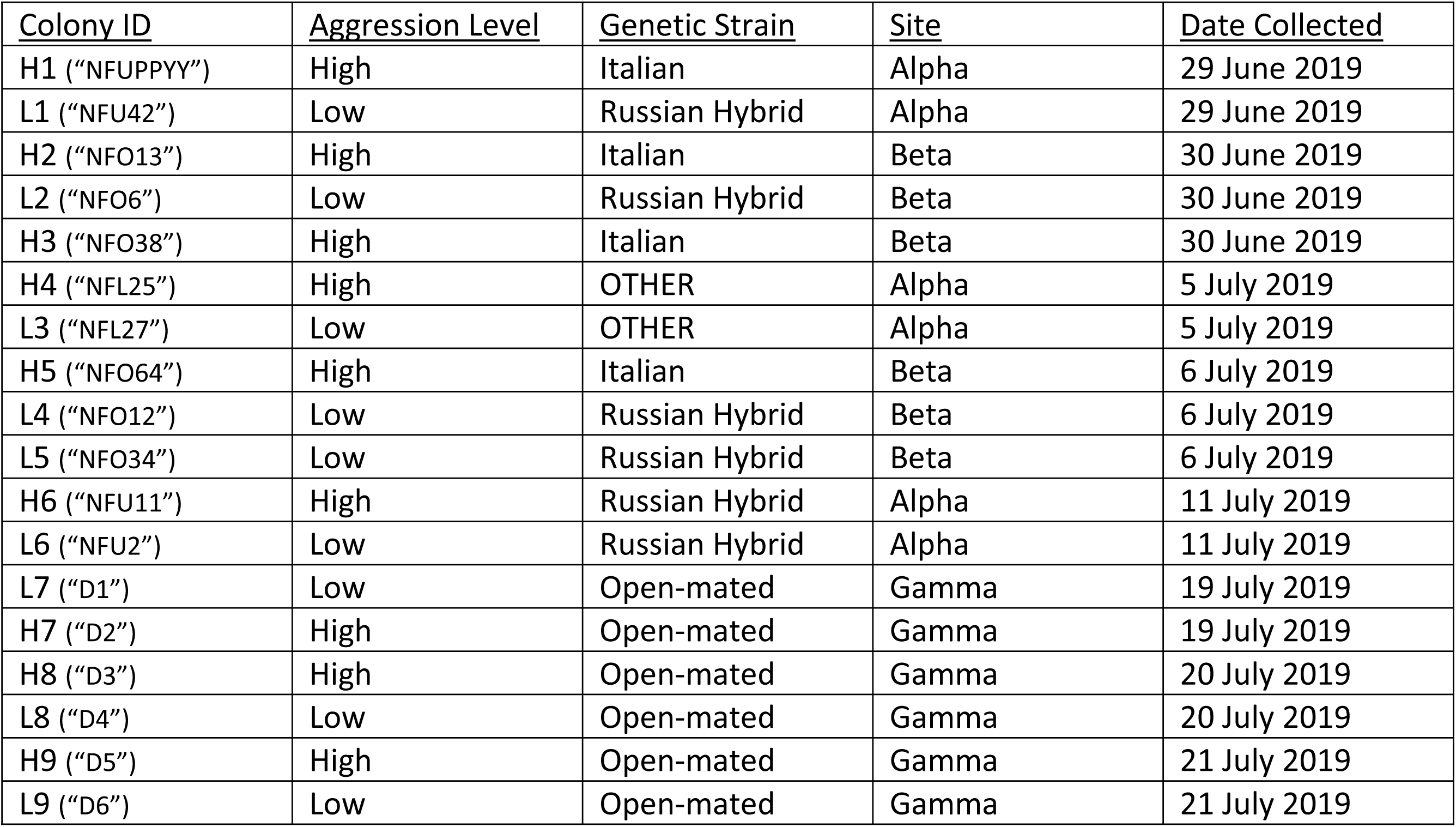
Table listing Colony ID, colony aggression level, genetic strain, site of the colony, and the date the worker jelly was collected.

**S11 Table.** Comparison of published studies examining the nutritional content of worker jelly. Proteins, lipids, and carbohydrates are reported as a mean percentage of the dry mass per sample (± standard deviation, where reported). Larval age (h) indicates the time post-egg hatching. This is typically a range because queens are given a range of time to lay eggs (see METHODS). In many cases, macronutrients per sample were simply reported as a mean. “Other” indicates the percentage of dry mass not accounted for by the three macronutrients (e.g. micronutrients, nondigestible material, etc.), and this is listed as a sample mean or a range depending on how results were reported in the study. “NR” indicates the information was not reported by the study.

| Study | Larval Age (h) | N (samples);<br>N (colonies) | Proteins<br>(% ± S.D.) | Lipids<br>(% ± S.D.) | Carbohydrates<br>(% ± S.D.) | Other<br>(%) | Total dry mass<br>(mg ± S.D.) |
| --- | --- | --- | --- | --- | --- | --- | --- |
| Planta 1889 | 0-96 | 2000; NR | 53.4 % | 8.4% | 18.1% | 20.1% | 1.7 |
| Kohler 1922 | 0-96 | NR; NR | NR | 23.3% | 15.7% | NR | NR |
| Haydak 1943 | 0-48 | NR | 78.3% | 17.7% | NR | NR | NR |
| Shuel & Dixon 1959 | 0-30 | 3-6; 2 | 49.2 ± 2.3% | 5.2 ± 1.1% | 12.0 ± 3.2% | 33.6% | NR |
| Brouwers et al. 1987 | 0-84 | 2-15; 3 | 40-65% | ~6-10% | 12-20% | 5-42% | NR |
| Kunert & Crailsheim 1988 | 0-96 | NR | 42-50% | NR | 9-19% | NR | NR |
| Wang et al. 2015 | 24-48 | 100; 5 | 50.7 ± 0.5% | NR | 9.6 ± 0.5% | NR | NR |
| **Present Study** | **24-48** | **467; 18** | **66.6 ± 40.1%** | **4.4 ± 2.1%** | **7.9 ± 4.8%** | **21.1 ± 27.1%** | **1.24 ± 0.63** |

**S12 Raw Data. Excel file with all raw data used to generate this manuscript.** This file has three sheets: the first sheet is a ReadMe, while the second and third sheets contain the two data files needed to generate all figures and statistical results reported in this study.

**S13 Code. PDF of an RMarkdown file displaying all code used to generate the figures and statistical results used in this study.**

